# GeTMoR: Simultaneous genomic, transcriptomic, and morphological profiling of rare single cells

**DOI:** 10.1101/2024.09.29.615279

**Authors:** Rishvanth K. Prabakar, Michael J. Schmidt, Peter Kuhn, James Hicks

## Abstract

Circulating tumor cells (CTCs), and circulating tumor related cells, are extremely rare cells that intravasate from the tumor into the circulatory system and can be captured via a liquid biopsy. Although CTCs contribute to the metastatic cascade, and diverse phenotypes of CTCs have been observed – including the cytokeratin expressing CTCs, CTC clusters, large polyploid CTCs, and CTCs undergoing epithelial to mesenchymal transition – little is known about their functionality. By virtue of CTCs being rare, a detection method that maximizes the information obtained per cell would be ideal to understand their biology and for use in diagnostic approaches. The challenge is that rare cell detection necessitates extensive processing steps, during which molecular content, such as RNA and DNA, needs to be preserved for downstream single cell analysis. We developed GEnomic, Transcriptomic, and MOrphological profiling of Rare cells (GeTMoR), a method that extends the High Definition Single Cell Assay for detecting rare cancer related cells to simultaneously image and profile the genome and transcriptome from single rare cells. We validated GeTMoR by spiking in cancer cell lines into whole blood to evaluate the quality of recovered gene expression and copy number profile from the same cell. The GeTMoR approach provides the ability to link the phenotype of rare cells, including CTCs, to their genome and transcriptome, thereby enabling insight into rare cell biology.

## 2 Introduction

Circulating tumor cells (CTCs) are extremely rare cells – typically detected at levels of 1 cell in 1 million immune cells [1] – that are recovered outside of the tumor environment through liquid biopsy approaches [2, 3]. Liquid biopsies in cancer research and care typically pertain to peripheral blood draws, but can also include sampling other bodily fluids, such as cerebrospinal fluid [4], bone marrow aspirates [5, 6], and portal vein sampling close to the primary tumor [7, 8]. CTCs are thought to play a crucial role in the complex process of seeding metastatic tumors from primary tumors [2, 9], and can thus be leveraged for interrogating tumor biology, diagnosis, and disease monitoring.

The CTC field has historically relied on studies that report on CTC abundance and morphological features [10]. These studies have shed light into the phenotypic diversity of rare cells in the bloodstream of cancer patients including a variety of cancer cell types distinguished by characteristically altered genomes [11, 12], such as epithelial CTCs, partial mesenchymal or complete mesenchymal CTCs [6, 13, 14], as well as platelet-coated CTCs [6], larger CTCs [15, 16, 17], CD45-positive CTCs [18, 19, 20], and apoptotic CTCs [21, 22]. While the genome and morphology of most of these cell types have been characterized, the transcriptional regulation of these cells types are unknown.

Additionally, rare non-epithelial tumor related cells that are not genomically altered and are presumed to arise from the tumor microenvironment (TME) have been defined. TME cells are known to play key roles in driving disease [23]. Recent reports have suggested that circulating TME (cTME) cells, such as cancer associated fibroblasts, have been detected in liquid biopsy samples of cancer patients [24, 25, 26]. A single-cell study in prostate cancer liquid biopsy highlighted CTC copy number clonality, but simultaneously revealed a substantial fraction of rare cancerrelated cells that were not copy number altered [11]. Profiling the rare circulating cell landscape of prostate and breast cancer liquid biopsy samples – including CTCs and putative cTME cells – a dramatic phenotypic diversity was identified in morphological characteristics through high definition immunofluorescence imaging and imaging mass cytometry [27]. Despite the presence of putative cTME cells, little is known about their functionality and relationship with disease.

The rarity of CTCs and tumor related cells necessitates using specific cellular properties that differentiate them from other blood cells for detection. These include selection via expression of epithelial proteins such as cytokeratin or EpCAM and the absence of leukocyte markers such as CD45, or through physical properties such as the larger size or deformability [28, 29, 30, 31]. While powerful, these techniques often miss subsets of CTCs and ignore unique cancer related cells, such as cTME cells. The High Definition Single Cell Assay (HDSCA) [13] is an imaging-based liquid biopsy system that detects rare cells by plating all nucleated cells on a microscopic slide, staining with markers to distinguish disease-related cells from millions of leukocytes, automated imaging and image analysis followed by picking these cells with a micro-manipulator for downstream single cell analysis. The HDSCA approach is optimized for rare cell detection – including CTCs and tumor related cells – and for genomic and proteomic analysis [11]. The benefit of the HDSCA is that it detects all cells without selective filtering and then determines rarity within that population. To date, molecular analysis has so far been limited to DNA copy number profiling and multiplexed proteomic profiling via imaging mass cytometry. Single cell transcriptional profiling would allow deeper understanding of the various cell phenotypes identified by imaging, however this has not been possible thus far on isolated rare cells using the HDSCA approach.

The challenge in performing any molecular analysis on CTCs and rare cells is that the required analytes need to be preserved during extensive protocols to detect and gain access to the cells. This is especially problematic for RNA which is known to be easily susceptible to RNAse degradation and leakage from permeabilized cells. Despite these challenges, several CTC detection methods have been used for single-cell RNA-seq analysis. Hydro-Seq [32], CTC-iChip [33, 34, 35], and the MagSweeper [36] have been used to isolate and subsequently profile the transcriptome from single CTCs. Further, CellSearch and Parasortix have been used to profile CTC expression signatures by isolating cells based on cell surface marker expression and morphology [37], respectively, but lack high resolution imaging of CTCs. These technologies are limited in only profiling a subset of cells selected by specific epithelial markers, such as EpCAM and cytokeratins, and miss other rare tumor related cells [14, 38].

Robust techniques are needed to further understand rare cells – including CTC and cTME cell – functionality. We re-engineered the HDSCA approach for characterizing tumor-related rare cells in circulation to perform transcriptomic analysis on the detected cells together with genomic and morphological characterization. We validated our approach by spiking cell lines into blood samples obtained from normal donors. We show that both the transcriptomic and genomic content from the same single cell are preserved after the steps to detect and isolate the spiked cells from millions of blood cells.

## 3 Results

The GeTMoR approach is conceptually based on the HDSCA approach for detecting tumor related rare cells in circulation [13]. The HDSCA approach depletes RBCs via ammonium chloride lysis, plating all nucleated cells on a specialized adhesion slide, followed by PFA fixation, immunofluorescence staining with cytoplasmic and membrane markers for WBC and tumor related cells (i.e., epithelial cell markers), imaging the entire slide, identifying potential rare cells, and preparing them for in-situ targeted proteomics or isolating them for downstream genomic analysis. The HDSCA approach is optimized for CTC detection and for genomic or proteomic analysis, however, the transcriptomic content of CTCs are degraded and lost through the process.

Molecular profiling of CTCs is inherently challenging given the rarity of these cells, and hence requires extensive processing steps to selectively capture CTCs. Moreover, profiling CTCs in a clinical setting necessitates the transport of blood samples from the point of collection to a laboratory for analysis, which typically takes 24 to 48 hours. In addition to these challenges, HDSCA fixes and stains for intracellular cytoplasmic markers (such as cytokeratins, vimentin, etc.) allowing for both higher sensitivity in detecting CTCs and for detecting a wider range of tumor related cells in circulation. Unfortunately, these factors make it challenging to profile the transcriptome from CTCs as RNA is highly susceptible to degradation.

### 3.1 Re-engineering HDSCA for transcriptomic profiling

We re-engineered HDSCA to address these limitations by optimizing each step of the process to preserve RNA while retaining the advantages offered by HDSCA. While we describe and evaluate the final version of GeTMoR in section 3.2, here we provide a rationale for the choices that were made for the critical steps of GeTMoR. A new user or a user adapting GeTMoR for their application will likely encounter similar challenges in preserving RNA. We provide this section in the hope that it will serve as a useful guide for method optimization.

#### Cleanliness and the use of RNAse inhibitor

RNAse is ubiquitous and extremely hard to inactivate. It only takes RNAse in any one step of the protocol for the entire protocol to fail. Therefore, it is of utmost importance that all equipment, pipettes, gloves, and lab bench be wiped down with RNAse inhibitor solution prior to start of experiment. It is also recommended to UV irradiate all reusable equipment such as pipettes, staining trays, wash jars, etc. Moreover, enzymatic RNAse inhibitor should be added fresh to every step of the protocol when applicable.

#### Extracting RNA and DNA from fixed single cells

The first challenge was to extract RNA from fixed cells that are plated on a microscopy slide. We adopted the FRISCR [39] protocol to extract both the DNA and RNA from the same cell by reverse cross-linking in the presence of heat and proteinase-K, followed by magnetic separation of RNA and DNA. Genomic DNA is recovered in the supernatant while RNA is bound to magnetic beads.

#### Blood collection tube and transportation time

Blood collected from patients is transported in a blood collection tube (BCT). Prior studies have evaluated different BCT types for the morphology, the DNA content, number of CTCs obtained [40, 41], and for the quality of RNA obtained [42] after 24 hours. RNA was well preserved in EDTA tubes for the GeTMoR protocol, and their ubiquitous availability made it an ideal choice. A GeTMoR user would have to evaluate several factors when selecting a BCT including availability at the collection site, the duration and temperature of transportation [40]. Ideally, blood samples should be processed as soon as possible after collection since cells are viable in an EDTA tube and could be prone to stress or cell state changes.

#### RBC depletion

The first step in processing a blood sample is to deplete RBCs. Typical approaches include ammonium chloride lysis, Ficoll-Paque separation, and antibody-based depletion [43]. The HDSCA approach uses ammonium chloride lysis, however, it was detrimental to RNA. Prior studies have shown that CTCs have a similar cell density to PBMCs, and separate out with the PBMC layer after centrifugation [44]. We thus evaluated Ficoll-Paque separation and we were able to extract RNA from spiked-in cancer cells that were obtained from the PBMC layer after separation.

#### Fixation

The cells plated on specialized adhesion slides are fixed with paraformaldehyde (PFA) to allow for cell permeabilization and intracellular antibody staining. Too little fixation could lead to loss of cytoplasmic RNA and proteins, and too much fixation may reduce the efficiency of reverse cross-linking required to extract RNA. We fix the cells with 2% PFA for 10 minutes.

#### Staining

The staining process has several points of failure that necessitated optimization, including the blocking buffer, permeabilization agent, and staining time. We found that RNA was better preserved in cells when using bovine serum albumin as opposed to goat serum, as RNAse content can vary dramatically in goat serum. Further, we found permeabilization with Saponin is less harsh on the cells as compared with detergents, such as Tween-20 or Triton-X, and hence was better at preserving RNA. Concentration of Saponin, permeabilization time, and staining time needs to be evaluated for each antibody. Using a higher concentration or longer time enables better antibody labeling, but also provides an opportunity for RNA degradation. The minimum concentration and time needed to achieve the optimal sensitivity for downstream application needs to be evaluated.

#### Slide scanning and image analysis

The longer the imaging and image analysis time, the more opportunity for RNA degradation. We minimized the imaging time by setting the microscope stage to move as fast as possible and by minimizing the exposure time. Moreover, we imaged the entire slide on one channel before switching to another channel to image the entire slide again. This greatly sped up the imaging due to not having to change imaging filters for each tile, but it comes at the expense of the channels being slightly misaligned (which was not an issue for detecting cells). We performed image analysis in real-time during imaging using a custom image analysis pipeline (described in supplementary section 1) written in C++ using the open-source ITK-Toolkit [45, 46].

### 3.2 GeTMoR for genomic, transcriptomic, and morphological characterization of rare cells

The GeTMoR (**Fig. 1**) protocol involves blood collection in an EDTA tube. The blood sample is processed using a standard Ficoll-Paque separation [47], and the mononuclear cells are extracted and washed. The cells are counted, and plated on an adhesion slide at approximately 3 million cells per slide. Multiple slides can be plated as needed. The cells are fixed with paraformaldehyde, permeabilized with Saponin, and stained with the nuclear marker DAPI, an epithelial marker pancytokeratin cocktail, mesenchymal marker Vimentin, and leukocyte marker CD45. The slide is loaded onto an IF microscope fitted with an automated stage. For detecting CTCs, whole slide imaging is performed only on the channels containing the nuclear and epithelial markers. The frame images are segmented to produce images of single cells. The segmented regions are used to obtain a feature vector consisting of the mean intensities, size, and shape of each cell. A frame consists mostly of WBCs with at most a few rare cells per frame. These rare cells are detected using the intensity of epithelial markers. The coordinates of these rare cells are used to locate them on the slide, which are then picked (and optionally imaged for additional markers at a higher resolution) from the slide into a PCR tube using a micro-manipulator. The isolated single cells can potentially be used for any downstream single cell analysis. We simultaneously extracted the genome from these cells for whole genome copy number profiling and transcriptome for full length mRNA profiling using an extended version of FRISCR [39]. GeTMoR could also be used in isolation either for only CTC detection, CTC copy number profiling, transcript profiling, or any combination of them. The detailed GeTMoR protocol is provided in supplementary section 2.

**Figure 1.**
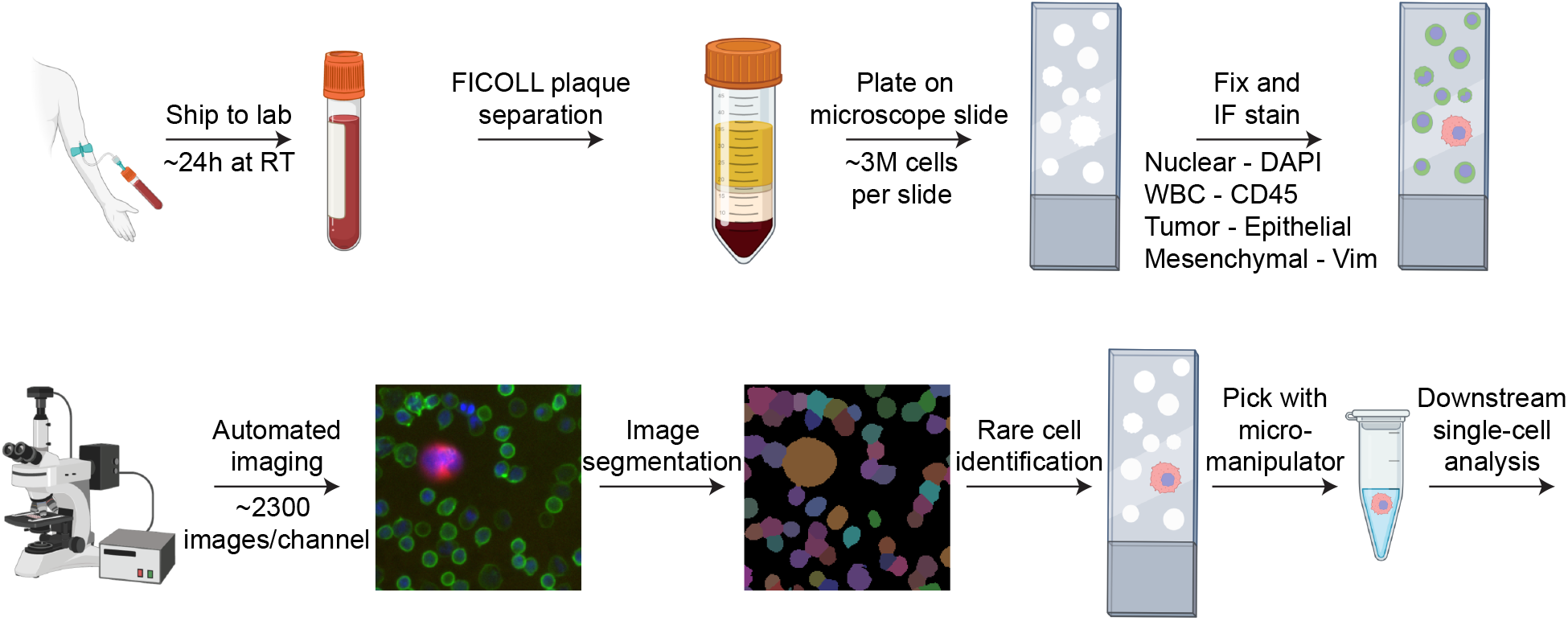
GeTMoR protocol for rare cell detection by extracting and plating all mononuclear cells, antibody labeling with WBC and tumor related markers, and automated imaging to detect rare tumor cells. These cells are picked for simultaneous single cell genomic and transcriptomic analysis. Figure created using Biorender.com

### Transcriptome and genome integrity of spiked-in cells

We validated GeTMoR by spiking cell lines into normal blood donor samples, performing all the steps of the GeTMoR protocol to detect and pick the spiked cells. High quality transcripts were recovered from spiked PC3 cells after GeTMoR protocol that were comparable to control PC3 cells (Extended figure 1). To evaluate the ability of GeTMoR to distinguish different cell types, a mix of approximately 1:4 MDA-MD-231 and PC3-GFP cells were spiked into normal blood samples obtained in EDTA tubes. We performed two replicates of this experiment. 35 cells were picked for genomics and transcriptomics in the first experiment and 11 cells were picked for the second experiment. The integrity of the genome and transcriptome of the spiked cells were determined by comparing them to control cell lines that were plated, fixed, and picked directly without going through the GeTMoR protocol (i.e., without being exposed to blood and subsequent staining and imaging steps).

To measure transcript integrity, the percentage of 5’ to 3’ coverage was measured for each condition, with increased 3’ coverage indicating degradation. We found that both spiked replicates displayed no substantial 3’ biases and were comparable with controls (**Fig. 2A**). As another quality metric, we measured the percentage of bases aligning to different genomic features and found they were comparable for controls and the spiked-in samples (**Fig. 2B**). We recovered at least 2,500 protein coding genes for each spike replicate, which provides sufficient information for downstream analysis (**Fig. 2C**). To classify spike-in samples as either MDA-MB-231 or PC3 cells, we determined the best genes to separate the two cell types using the control cells with SingleR [48]. We then scored each spike-in cell and clustered the cells based on the reference genes. The classified spike-in cells cluster together with the respective controls. (**Fig. 2D**). Taken together, this data shows GeTMoR processing retains transcript quality in spiked-in cells, and allows for cell type classification compared to a control ground truth data set.

**Figure 2.**
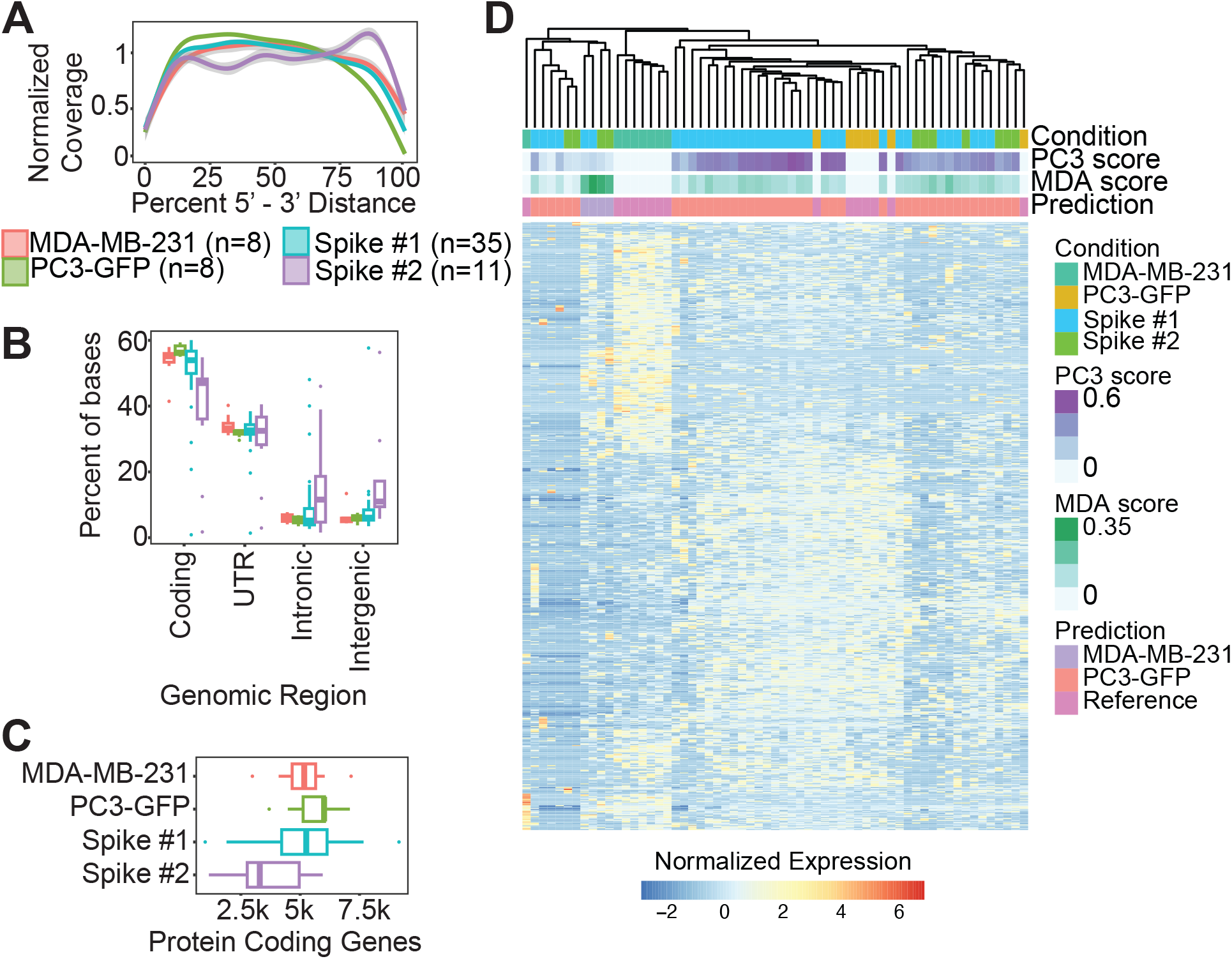
Recovered transcript quality of GeTMoR processed cells. (A) Normalized transcript coverage from 5’ to 3’ end for each condition. 3’ accumulation indicates degradation. Color key refers to all panels in the figure. (B) Percentage of bases mapped to distinct genomic features. (C) Total expressed protein coding genes for control MDA-MB-231 and PC3 samples, as well as GeTMoR-processed replicates 1 and 2. (D) SingleR marker analysis for all samples using identified reference genes between PC3 and MDA-MB-231 control samples.

**Figure 3.**
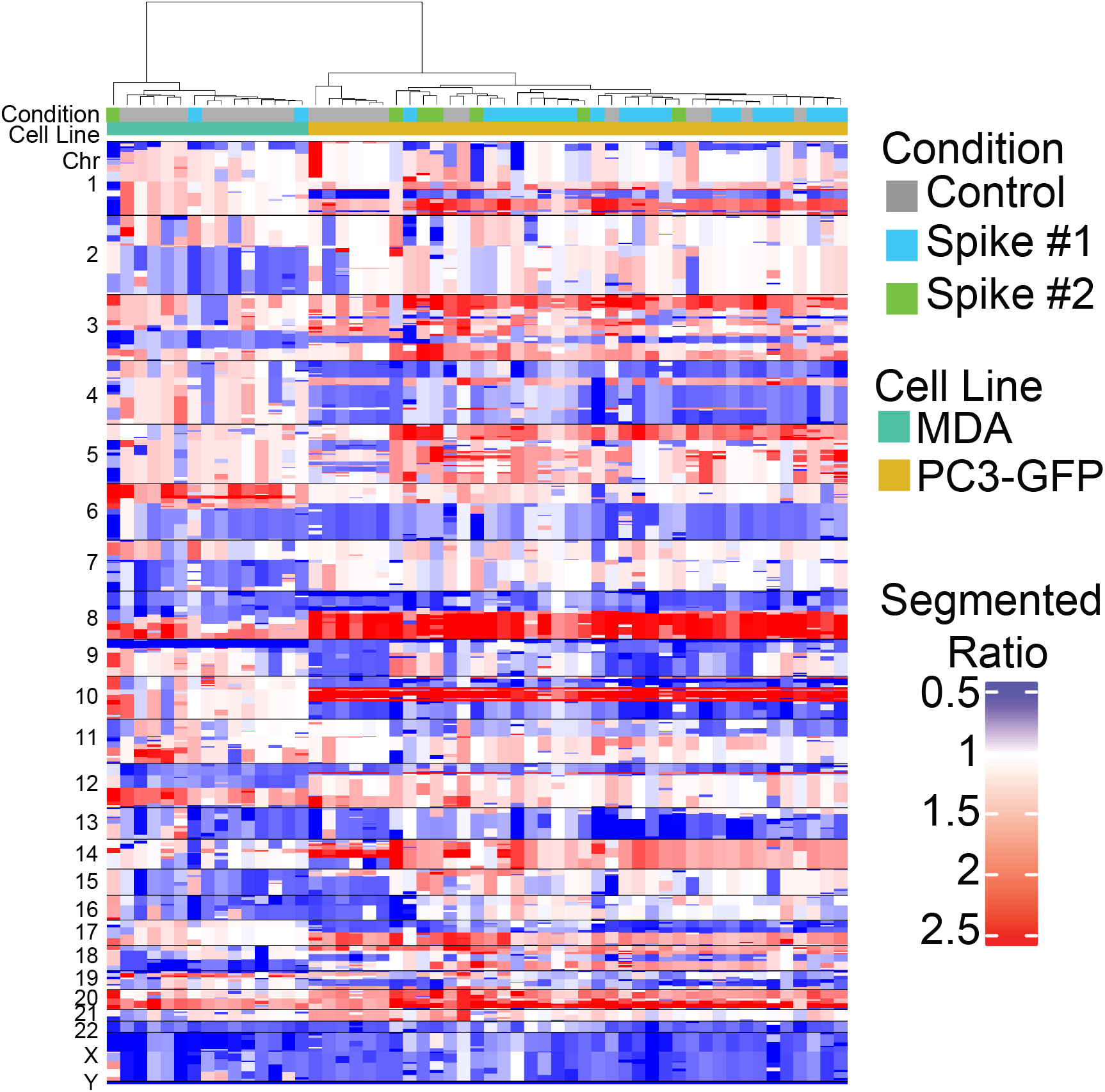
GeTMoR retains DNA copy number profiles for MDA-MB-231 and PC3 spiked cell lines. Single cell segmented ratio copy number profiles from same single cells presented in Figure 2.

**Figure 4.**
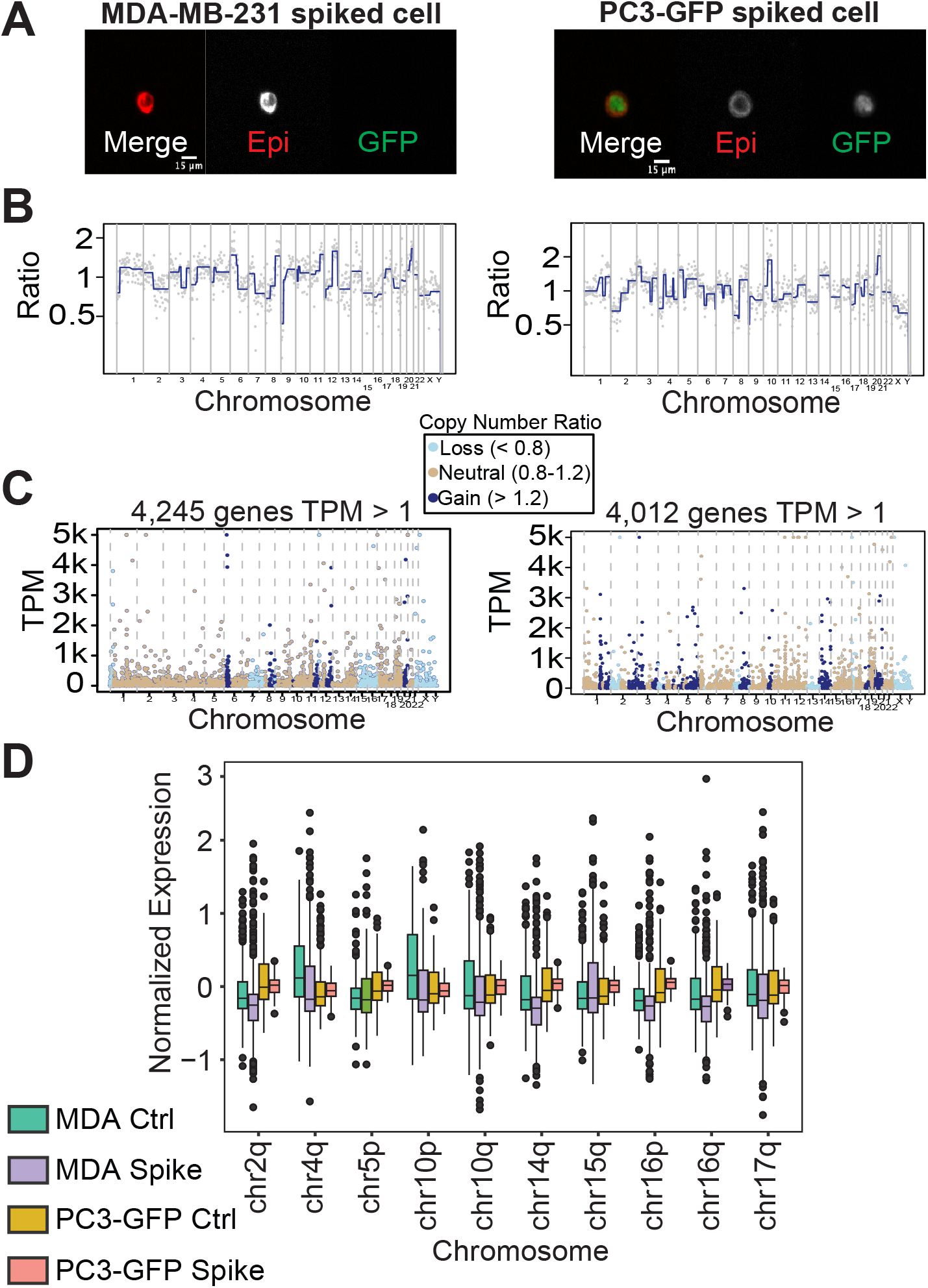
Simultaneous single cell profiling of morphology, genome, and transcriptome. (A) Representative image of MDA-MB-231 and PC3 spiked-in cells processed with GeTMoR (B) Copy number profiles from cells isolated in (*A*). (C) Gene expression (TPM) profiles plotted against chromosomal location and copy number ratio for cells in (*A*). (D) Chromosomal arm versus gene expression for MDA-MB-231 and PC3 control and spiked cells

We then compared the copy number profiles obtained from the same cells to the control cell lines. As expected, the PC3 and MDA-MB-231 cells clustered with the respective controls after GeTMoR processing (**Fig. 3**).

### Simultaneous genomic, transcriptomic, and morphological profiling of single cells

The complementary components of GeTMoR enable unprecedented insight into single cells. As a proof of principle, we used PC3-GFP spiked cells to distinguish MDA-MB-231 from PC3 cells through fluorescence imaging. Through image analysis, PC3 cells were identified via GFP positivity, while MDA-MB-231 cells were negative for GFP (**Fig. 4A**). Copy number ratio data further distinguished MDA-MB-231 from PC3 cells and confirmed by image analysis (**Fig. 4B**). To visualize how copy number can influence gene expression, we plotted chromosomal position by TPM for individual cells (**Fig. 4C**). Lastly, we compared the gene expression (aggregated for all cells in each category) of chromosomal arm regions with differing copy numbers (**Fig. 3** and **Fig. 4D**). Copy number neutral and gained regions typically have higher gene expression values than genomic regions with fewer chromosomal copies.

## 4 Discussion

The aim of GeTMoR is to provide a robust method that enables the transcriptomic profiling of CTCs and other rare cells. GeTMoR approach offers several advantages over other methods for isolating rare cells for molecular analysis: (1) GeTMoR can extract both RNA and DNA from the same rare cell of interest, and the approach can potentially be extended to other assays such as CITE-seq for protein expression analysis [49] or bisulfite sequencing for methylation profiling [50, 51]. Further, GeTMoR may also be applicable to existing technologies that detect CTCs and other rare cells through imaging, such as the RareCyte CTC detection system which is readily used in many CTC studies [52]. (2) The assay allows for fixing and permeabilizing cells and thus can stain for cytoplasmic proteins for protein detection without substantial transcript loss. This enables interrogating a wider variety of CTCs and tumor related cells such as platelet coated CTCs, epithelial to mesenchymal transitioning CTCs, and cancer associated fibroblasts. (3) GeTMoR provides simultaneous views of the phenotype (IF image), cell state (RNA), and genotype (DNA) for each cell, thus enabling a deeper insight into rare cells.

There are multiple benefits to GeTMoR, but some limitations exist. RNA degradation occurs rapidly during processing, thus optimal use of the GeTMoR protocol requires that blood processing, slide preparation, imaging and single cell isolation be performed on the same day the sample is received. While improvements have been made to GeTMoR to enhance the chances of successfully capturing intact transcripts, there will inevitably be some cells that do not pass RNA-seq quality control due to transcript degradation. Further, to expedite the time required before single cells are isolated and minimize transcript degradation, GeTMoR only images in one or two fluorescent channels. While one to two channels still provides detailed morphological data, we acknowledge the user may want to visualize more protein markers especially in the context of cTME cells. Imaging additional channels will require further method optimization for the user specific purposes (eg., image a smaller area of a slide). Lastly, we have not estimated the sensitivity and specificity of GeTMoR to detect CTCs. While this is not crucial when using GeTMoR to interrogate the biology of CTCs, but it is crucial if the number of detected CTCs is used as a metric in any downstream analysis.

While studying the transcriptome of CTCs and other rare cells is challenging, GeTMoR offers a robust technique to obtain as much information as possible from a single cell. Capturing the genomic content to understand clonal lineage, transcriptome to understand the state of the cell, and morphology to visualize cellular size and texture allows for unprecedented multi-modal analysis that has the potential to advance CTC biology.

## Methods

### Cell line collection and blood processing

PC3 and MDA-MB-231 cells were purchased from ATCC (Manassas, Virginia) and grown in RPMI and DMEM, respectively, with 10% FBS and 0.5% penicillin/streptatvidin (Sigma, P4333). GFP-PC3 was purchased from Angioproteome and grown in RPMI (cAP-0067GFP). Cells were seeded two days prior to each experiment in a T-25 flask and collected at roughly 85% confluence. Cells were then lifted with 1x versene solution, spun down at 1000 RPM for 5 minutes, and resuspended in 1x PBS with RNAse inhibitor. Blood from normal donors (no disease) was diluted 1:1 in 1x PBS with RNAse inhibitor. Cell lines were then spiked into the diluted blood. The blood mixture was then added to 15 milliliters of Ficoll and spun down at 400xG for 20 minutes. Nucleated cells at the top of the Ficoll layer were recovered and washed in 1x PBS with RNAse inhibitor and then spun down at 1000 RPM for 5 minutes. Cells were then plated on specialized Marienfield glass slides (Marienfeld, Lauda, Germany) and incubated at 37^*°*^C for 15 minutes.

### Immunofluorescence staining

Slides were removed from the incubator and immediately fixed with 2% paraformaldehyde for 10 minutes. Slides were blocked with 2% BSA in PBS and 4,6-diamidino-2-phenylindole (DAPI; D1306, Thermo Fisher Scientific, Waltham, MA, USA) for 5 minutes and permeabilized with 0.1% Saponion (Thermo, A18820-22). Then, slides were incubated at room temperature with a primary antibody cocktail consisting of mouse IgG1/Ig2a antihuman cytokeratins (CK) 1, 4, 5, 6, 8, 10, 13, 18, and 19 (clones: C-11, PCK-26, CY-90, KS-1A3, M20, A53-B/A2, C2562, Sigma, St. Louis, MO, USA), mouse IgG1 anti-human CK 19 (clone: RCK108, GA61561-2, Dako, Carpinteria, CA, USA), and mouse EpCAM. The CK and EpCAM antibodies make up the EPI channel. Conjugated mouse anti-human CD45 Alexa Fluor 647 (clone: F10-89-4, MCA87A647, AbD Serotec, Raleigh, NC, USA), rabbit IgG anti-human vimentin (Vim or V) (clone: D21H3, 9854BC, Cell Signaling, Danvers, MA, USA), and DAPI were also added to the primary incubation buffers. Slides were then washed in 2% BSA in PBS and incubated at room temperature for 10 minutes with Alexa Fluor 555 goat anti-mouse IgG1 antibody (A21127, Invitrogen, Carlsbad, CA, USA), conjugated mouse anti-human CD45 Alexa Fluor 647, VIM, and DAPI. Slides were washed, cover-slipped, then immediately scanned.

### High content scanning and image analysis

Automated high-throughput fluorescence scanning microscopy was performed in the TRITC (CK) and FITC (GFP) channels with a 10x objective lens, collecting 2304 frames per channel.

Images generated from each channel were smoothed, thresholded to convert the grayscale image to binary cell consisting of only the cellular regions in the foreground, and individual cells were labeled using connected components. Cell coordinates and features were extracted from the labeled cells. Optionally, cell masks from multiple channels can be merged before labeling individual cells using watershed segmentation. The image analysis pipeline was implemented using the ITK Toolkit [46]. The image analysis pipeline is described in detain in Supplementary section 1.

### Single cell isolation

Rare cells were identified by X and Y coordinates on the slide and isolated from slides using a robotic micromanipulator system (Eppendorf). Individual cells were deposited into PCR strips containing 1x PBS with 0.2% Triton X-100 and RNAse inhibitor. Cells were then stored on dry ice and then placed in the -80*°*C until RNA and DNA extraction.

### DNA and RNA extraction

RNA and DNA were first reverse cross-linked via a modified FRISCR approach [39]. Briefly, lysis buffer was added to the cell (0.2% Triton X-100 + RNase inhibitor) and incubated at 56*°*C for 1 hour. dT25 beads (NEB, S1419S) were then added to the samples and hybridized at 56*°*C for 1 minute. Samples were incubated at room temperature for 10 minutes then added to a magnetic PCR rack. DNA was collected in the supernatant while RNA remained attached to the beads.

DNA was removed in the supernatant and was subsequently amplified with single cell whole genome amplification (WGA; Sigma-Aldrich; Cat# WGA4). Libraries were constructed using the DNA Ultra FSII Library Prep Kit (New England Biolabs; Cat# E7430). Cells were sequenced 150 base-pairs paired end on an Illumina HiSeq 4000 (Fulgent) at a depth of 1-2 million reads per sample.

RNA was washed once with 2x SSPE buffer and once with 1x PBS, and then eluted in RNAse free water at 80*°*C. Purified RNA then underwent a modified Smart-Seq2 protocol. Briefly, dT primer was added to the purified RNA and allowed to hybridize for 3 minutes at 72*°*C. Reverse transcription and template switching then took place at 42*°*C for 90 minutes, followed by 10 rounds of cycling from 50*°*C for 2 minutes and 42*°*C for 2 minutes. cDNA was then amplified and sequencing libraries were prepared with Nextera XT (Illumina). Cells were sequenced 150 base-pairs paired end on an Illumina HiSeq 4000 (Fulgent) or an Illumina NovaSeq (Novogene) at a depth of 1-2 million reads per sample.

### Single cell RNA-seq analysis

Adapters were trimmed using TrimGalore [53] (version 0.6.10) with the parameters `--paired --quality 20 --stringency 3`. Trimmed reads were aligned to reference genome hg38 and Gencode annotation version 44 using STAR [54] (version 2.7.11) with the following parameters --readFilesCommand zcat --soloType SmartSeq --soloUMIdedup Exact NoDedup --soloStrand Unstranded --outSAMattributes RG NH HI nM AS --outSAMtype BAM SortedByCoordinate --outFilterScoreMin 30 --soloFeatures GeneFull. Picard (version 3.2.0) was used to visualize RNA mapping quality control. Seurat [55] (version 5.0.3) and SingleR [48] (version 1.10) were used for comparison with controls and cell type classification.

### Single cell copy number profiling

Copy number profiling from low pass whole genome sequencing samples was conducted as previously described [56, 57]. Briefly, reads were aligned with BWA-MEM [58] to the hg38 reference. PCR duplicates were removed and counts were determined for 5000 bins containing a equal number of uniquely mappable locations. Count data was normalized for GC content and segmented via the R package DNACopy (version 1.70.0).

## Code and data availability

Image analysis code used for GeTMor is available at https://github.com/CSI-Cancer/getmor_segmentation. All data is available upon request from the authors.

## Author Contribution

**Conceptualization**: RKP, MJS, JH; **Methodology**: RKP, MJS, JH; **Software**: RKP; **Validation:** RKP, MJS; **Formal analysis**: RKP, MJS; **Investigation**: RKP, MJS; **Resources**: PK, JH; **Writing -Original Draft**: RKP, MJS; **Writing -Review & Editing**: RKP, MJS, PJ, JH; **Visualization**: RKP, MJS; **Supervision**: JH; **Funding acquisition**: PK, JH.

## Acknowledgments

We would like to thank the lab members at CSI-Cancer, particularly Amin Naghdloo, Nikki Higa, and Mohamed Saleh for their support throughout the development of GeT-MoR.

## Supplementary Figures

**Extended figure 1.**
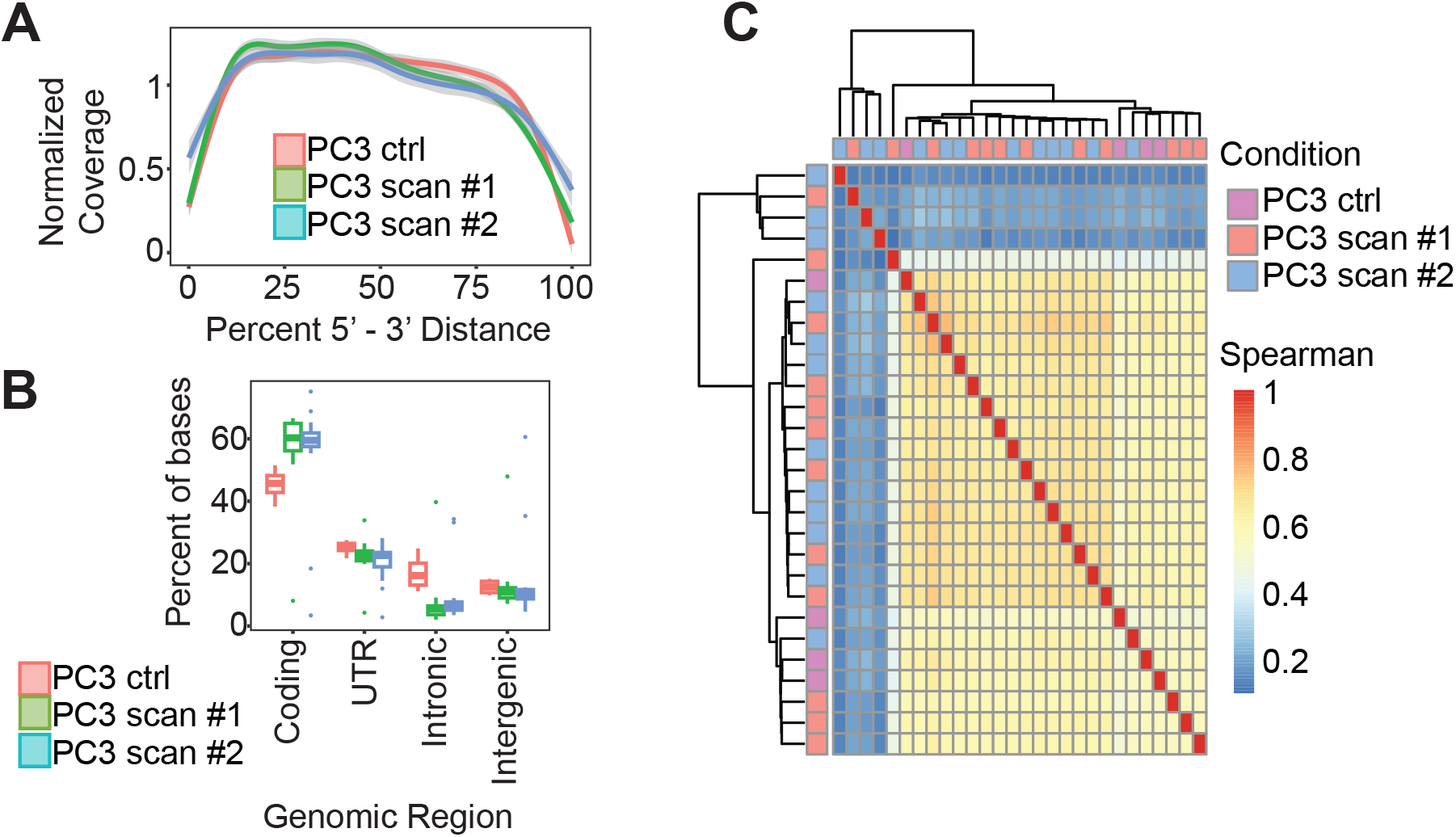
Recovered transcript quality of spiked PC3 cells after GeTMoR protocol is comparable to control PC3 cells. (A) Normalized transcript coverage from 5’ to 3’ end for each condition. 3’ accumulation indicates degradation. Color key refers to all panels in the figure. (B) Percentage of bases mapped to genomic regions. (C) Spearman’s correlation of gene counts.

