## Supplementary text for "GeTMoR: Simultaneous genomic, transcriptomic, and morphological profiling of rare single cells"

### 1 Supplementary notes

#### 1.1 Image segmentation and rare cell detection

The GeTMoR approach detects rare cells on a slide through automated scanning of the entire slide. All mononuclear cells (obtained after Ficoll separation), including lymphocytes, monocytes, and potential circulating tumor cells (CTCs), are plated on the slide. The slide is stained with a combination of markers to differentiate CTCs from leukocytes. It is then imaged across multiple channels using an automated scanning microscope. The scanned images are processed to detect the number and location of CTCs on the slide, which can subsequently be isolated for downstream single-cell analysis.

The automated scanner images the entire slide generating thousands of frames per slide, and one image per frame for each channel imaged. A slide typically consists of $\sim$3 million cells almost all of which are leukocytes. CTCs, if present, are typically in the range of $\sim$1-10 cells per slide. The objective of the image analysis pipeline is to process the images to (1) segment the cells from the slide background and (2) detect the CTCs from all the cell images on a slide.

We implemented an image segmentation pipleine using the ITK toolkit. The pipeline was designed with the following considerations:

1. Speed: A crucial aspect of preserving the RNA in cells is to minimize the time from when the cell is plated on a slide to when a cell is picked from a slide into a tube for long-term storage. We designed the computational image analysis pipeline to process the images in parallel with the slide scanning, and hence cells can be picked right after scanning.

2. Ability to process any number of channels: The slide can be scanned on any number of channels depending on the application. At minimum, a slide is scanned on the channel stained for CTC specific maker to detect the cells of interest. The maximum number of channels scanned is limited by the scanning microscope (typically 4-6 channels). We designed our pipeline to work irrespective of the number of scanned channels.

A slide typically consists of two categories of cells: abundant cell types such as WBCs and rare cell types such as CTCs. The images generated for makers for common cell types have different properties than images generated for markers for rare cell types. As an example, consider the images of WBC maker CD45 and CTC maker pan-cytokeratin (CK). Since almost all cells on a slide are WBCs, all frames will contain cells that are positive for CD45 (typically $\sim$1500 cell per frame). Whereas, since CTCs are rare, almost all frames will have no signal on the CK image and frames that have a positive signal will contain at most a few cells (typically 1-3 cells per frame). The image segmentation steps for these two frame types differ. The image segmentation pipeline consists of the following steps:

**Image smoothing:** The images are blurred with a Gaussian anisotropic diffusion image filter that preserves the edges of cells while smoothing the image.

**Cellular fraction estimation:** An approximate fraction of pixels in frames that are expected to contain cellular foreground pixels is used in the next step to determine a threshold for segmentation. An approximate cellular fraction can be directly specified as an user parameter. As an example for frames containing only rare cells most pixels would belong to the background and so a high value such as 99.5% can be used. Alternatively, the cellular fraction can be estimated by thresholding the input frame and using the fraction of foreground pixels as the cellular fraction.

**Determining cellular foreground pixels:** The cellular foreground pixels are determined using a double thresholding approach that uses a low and high threshold. The low threshold is a lenient threshold selected such that all the pixels that are a part of a cell are in the foreground, but at the expense of having non-cellular foreground pixels. The high threshold is a stringent threshold selected such that at least one pixel in a cell is in the foreground, and no pixels from the non-cellular components. The low thresholded image is used as a mask and the high thresholded image is used as a seed, and the double thresholded image is constructed by selecting all the foreground pixels in the lower threshold image that are connected components of the foreground pixels in the high thresholded image using the process of reconstruction by dilation.

The low threshold is determined by computing the approximate fraction of the frame that are covered with cells, and using the quantile intensity of the frame at $(1 - \mathrm{cell\_fraction} + \mathrm{offset})$ as the threshold. Optionally, an offset can be added to the cell fraction before computing the quantile. A positive offset would increase the quantile, and is useful for certain markers (such as CK or Vimentin) where not all cells in the frame are expected to express these markers. As an example, the cellular fraction for a typical frame is 0.15 and the lower threshold for each channel would be the 85th quantile of that channel with an offset of 0, or 90th quantile with an offset of 0.05.

Two different approaches are used to determine the upper threshold depending on the channel. For channels that are always expected to have a high number of cells (such as DAPI and and CD45), the upper threshold is calculated based on the quantile of the foreground pixels from the lower thresholded image. For example, a quantile of 0.5 would use the median intensity of all the foreground pixels in the lower thresholded image as the upper threshold. This approach would not work for channels that may not have a cell such as CK. If the CK channel does not have a true cell, a quantile, however high, would always result in a false positive seed in the upper thresholded image. For such channels, the upper threshold quantile is calculated as the smallest quantile that is at least a user specified multiple of the median. The upper threshold is then the intensity of the foreground pixels at that quantile. If no pixels satisfy this criterion, the upper threshold is set to infinity, and the entire frame is to background. This ensures that a channel will not have any false positives if it does not have a cell.

**Merging cellular foreground across channels:** Optionally, the individual channel masks are summed together to generate one frame mask. This frame mask contains a foreground pixel at locations containing foreground pixels in any of the individual channel masks. Alternatively, this step can be skipped and each channel can be segmented individually.

**Segmenting individual cells:** Two different segmentation procedures are used depending on if the frames contain rare cells or abundant cells.

For frames that contain rare cells, the probability of two cells touching one another are very low. Connected components are determined and each component is labeled as a cell.

For frames that contain abundant cell types, there would almost certainly be touching cells. These touching cells should be labeled as two distinct cells. The binary mask is distance transformed and a watershed segmentation is applied to label individual cells.

**Feature extraction:** The following shape features are extracted for each labeled cell: the cell x- and y- coordinates, cell area (in pixels), cell elongation, cell roundness. The following intensity features are extracted for each labeled cell for each channel: mean intensity, standard deviation of intensity, minimum intensity and maximum intensity.

### 2 GeTMoR protocol

#### 2.1 Protocol considerations and notes

- GeTMoR protocol performed on the same day as the blood draw is always preferred and will work the best; the sooner the better so the cell do not change expression programs while in the blood tube, and does not degrade.
- GetMoR protocol takes approximately 3 hours and should be performed as soon as a blood sample is received.
  - Blood processing and slide preparation takes $\sim$90min.
  - Fixing and staining takes $\sim$45min.
  - Scanning and finding cells of interest takes $\sim$30min.
  - Picking rare cells of interest will vary depending on the number of cells isolated.

  Once single cells are isolated, they can be stored at -80°C long term.

#### 2.2 Detailed GeTMoR protocol:

**Equipment:**

- Microwave
- Centrifuge (15mL and 50mL tube racks)
- Automated scanner or imaging apparatus
- Picking apparatus for single cell isolation
- RNAse-free laminar flow hood
- 96-well magnetic plate
- 96-well cooler plate
- Filtered pipette tips 1μL, 200μL, 1000μL

- Serological pipette
- Thermocycler
- UV hood for sterilization

**Reagents:**

- Nuclease free water (Invitrogen: 10977-015)
- 10x PBS (Invitrogen: 70011-044)
- EDTA (Sigma: 03690)
- RNAse inhibitor (NEB: M0314L)
- RNAse Away (VWR: 7003)
- RNAse free BSA (Sigma: 126615)
- Ficoll paque premium (VWR: 17544202)
- Saponin (Sigma: 47036)
- dT25 magnetic beads (NEB: S1419S)
- 20x SSPE hybridization buffer (Sigma: S2015-1L)
- Mouse IgG1/IgG2a anti-human cytokeratin (CK) 1, 4, 5, 6, 8, 10, 13, 18, and 19 (clones: C-11, PCK-26, CY-90, KS-1A3, M20, A53-B/A2, C2562, Sigma)
- Mouse IgG1 anti-human CK 19 (clone: RCK108, GA61561–2, Dako)
- CD326 (EpCAM) Monoclonal Antibody (1B7), eBioscience (Thermo Fisher Scientific: 14-9326-82)
- Mouse antihuman CD45:Alexa Fluor 647 (clone: F10–89–4, MCA87A647, AbD Serotec)
- Rabbit IgG antihuman vimentin (VIM): Alexa Fluor 488 (clone: D21H3, 9854BC, Cell Signaling Technology)
- Alexa Fluor 555 goat anti-mouse IgG1 antibody (Invitrogen: A21127)
- 4,6-diamidino-2-phenylindole (DAPI; D1306, Thermo)
- Triton-X 100 (Thermo: 327372500)
- 50% Tween 20 (Life Technologies: 003005)
- Reverse transcriptase (Takara: 639537)
- dNTP mix (NEB: N0447S)
- dT primer: (5–AAGCAGTGGTATCAACGCAGAGTACT(30)VN-3) ordered from IDT

- TSO primer: (5-AAGCAGTGGTATCAACGCAGAGTACATrGrG+G-3) ordered from IDT
- PCR primer: (5-AAGCAGTGGTATCAACGCAGAGT-3) ordered from IDT
- KAPA 2x HiFi mix (Rosche: KK2602)
- Nextera XT library prep kit (Illumina: Nextera XT library prep ki)
- Single cell DNA amplification kit (Sigma: WGA4)
- NEB Ultra FS II library prep kit (NEB: E6177)
- Marienfield glass slides (custom adhesive coating)
- HybriWell sealing glass coverslips (Grace BioLabs: HBW2260FL)

**Preparation:**

1. Wipe down all bench surfaces, pipettes, wash jars, and equipment with RNAse away. If possible, UV irradiate all wash jars and pipettes. Rinse all wash jars with nuclease free PBS.
2. 2% PFA preparation: Add $3.5\text{mL}$ of 1x PBS in a Falcon tube and add $0.5\text{mL}$ of 16% PFA. This can be made when preparing staining buffers during the 20 minute spin down of FICOLL.
3. Prepare stock of 1% saponin and store in 4°C. Good for 1 month

**Blood processing:**

1. Prepare the processing buffer: To $50\text{mL}$ of 1x PBS in a $50\text{mL}$ Falcon tube, add 100µL of 0.5M EDTA.
2. Add $15\text{mL}$ of FICOLL into a separate $50\text{mL}$ Falcon tube.
3. Take a $50\text{mL}$ Falcon tube and add processing buffer equal in volume to the amount of blood sample to be processed. For every $1\text{mL}$ of processing buffer, add $0.5\mu\text{L}$ of RNAse inhibitor. Gently add an equal volume of blood to the tube. Mix by inverting the tube.
   (a) $2\text{mL}$ of blood are used to make 1 slide. Therefore, in most cases, you should add $2\text{mL}$ of blood, $2\text{mL}$ of 1x PBS, and 1µL of RNAse inhibitor.
4. Very gently pipette the diluted blood onto the FICOLL, being careful not to mix the layers.
5. Centrifuge at 400xG for $20\text{min}$ at room temperature with acceleration set to 3 and deceleration set to 0.
6. During the centrifugation, prepare slides for plating as described in the cell plating section. Also, prepare the blocking, primary, and secondary staining solutions as described in the staining section.
7. At the end of the centrifugation, add $3\text{mL}$ of processing buffer to a $15\text{mL}$ Falcon tube and add 3µL of RNAse inhibitor.
8. Using the $15\text{mL}$ Falcon tube, carefully pipette the entire PBMC layer into the processing buffer, being careful not to pipette any FICOLL and avoiding the plasma layer. Mix by gently inverting the tube.

9. Centrifuge at 1000 RPM for 5min at room temperature with acceleration and deceleration set to 3.

10. Pipette out and discard the supernatant, being careful not to disturb the cell pellet.

11. Resuspend the cells in 1mL of processing buffer (with RNAse inhibitor) per slide and gently mix by pipetting. The required volume depends on the plating density of the slides and the cell count. If you are processing 1 slide and using 2mL of blood, then add 1mL of processing buffer.

**Cell plating:**

1. (Steps 1 to 4 can be performed during the FICOLL centrifugation step) Add the slide to a glass Coplin jar. Add RNAse-free water to a wash jar and add 4μL of RNAse inhibitor. Microwave for 30s.

2. Add 1x PBS to a wash jar and add 4μL of RNAse inhibitor.

3. Transfer the slides to the wash jar containing PBS.

4. Remove excess PBS from the slide and add 750μL of resuspended cells onto the slide.

5. Incubate at 37°C for 15min to allow the cells to adhere to the slide.

6. Remove excess buffer from the slide.

**Fixing and staining:** The goal of this staining protocol is to antibody label the rare cells of interest. This staining protocol can be modified with the appropriate antibody panel.

1. Prepare 2% PFA. Add 1mL of 2% PFA to the slide and let the cells fix for 10min at room temperature.

2. Prepare 2x PBS DAPI: Add 5mL of 10x PBS to 20mL of RNAse-free water to make 2x PBS. Then add 1μL of DAPI. Mix and use as the base buffer for the staining solutions.

3. Blocking solution preparation: 500μL of 2x PBS-DAPI, 100μL of 10% BSA, 396μL of nuclease-free water, 4μL of RNAse inhibitor, and 10μL of 1% Saponin.

4. Primary antibody solution preparation: 500μL of 2x PBS-DAPI, 100μL of 10% BSA, 361μL of nuclease-free water, 4μL of RNAse inhibitor, 10μL of primary pan-CK antibody, 5μL of primary CK-19 antibody, 10μL of EpCAM, 10μL of CD45-Alexa-647 conjugated antibody.

5. Secondary antibody solution preparation: 500μL of 2x PBS-DAPI, 100μL of 10% BSA, 372μL of nuclease-free water, 4μL of RNAse inhibitor, 10μL of CD45-Alexa-647 conjugated antibody, 10μL of Vimentin-Alexa-488 conjugated antibody, and 4μL of Mouse-IgG-Alexa-555 antibody.

6. Add 1x PBS to a wash jar and add 4μL of RNAse inhibitor.

7. Remove PFA from the slide, transfer the slide to the wash jar, and wait 30s.

8. Add 1mL of blocking solution to the slide and wait 5min.

9. Remove the blocking solution from the slide.

10. Add 1mL of primary antibody solution to the slide and wait 10min.

11. While waiting, prepare the automated image scanner. Be sure to wipe down the stage and the scanner with RNAse Away.

12. Add 1x PBS to a wash jar and add 4μL of RNAse inhibitor.

13. Remove the primary antibody solution, transfer the slide to the wash jar, and wait 30s.

14. Add 1mL of secondary antibody solution to the slide and wait 10min.

15. Mounting media preparation: To 200μL of 1x PBS, add 0.4μL of EDTA, and 4μL of RNAse inhibitor.

16. Add 1x PBS to a wash jar and add 4μL of RNAse inhibitor.

17. Remove the secondary antibody solution, transfer the slide to the wash jar, and wait 30s.

18. Remove excess PBS from the slide and add 200μL of mounting media.

19. Coverslip with RNAse-free HybriWell coverslips.

20. Prepare picking buffer by taking 10x lysis buffer (2% Triton X-100) and making a 1x working solution (1:10 dilution). Generally, making 200μL is sufficient (180μL of RNAse-free water + 20μL of 2% Triton X-100). Place on ice until ready for use.

21. Prepare slide buffer by adding 1μL of 50% Tween-20 to 10mL of 1x PBS. Vortex vigorously for at least 10 seconds to mix. Place on ice until ready for use.

**Scanning:** The scanning protocol will vary depending on the make and model of the automated scanning microscope. We describe a general scanning procedure without specific details. For faster scanning, the entire slide can be image on one channel before image the entire slide on the next channel (as opposed to imaging the all channels for one frame before moving to the next). By imaging on the channels containing rare cells first, image analysis can be performed in parallel while imaging.

1. Wipe down the stage and scanner with RNAseAway.

2. Open the scanner application and perform any scanner initialization and calibration.

3. Gently place the slide on the stage.

4. Select channels to be imaged and set the exposure time and gain.

5. Create support points for tile imaging and set the focus at each point using the DAPI channel.

6. Fix a fiducial point on the slide, and note the stage coordinates at the fiducial point.

7. Scan image the slide. Keep track of the stage coordinates for each frame to be able to re-locate any cell for downstream cell picking.

**Image segmentation and rare cell detection:** The parameters for the programs below are based on the assumption that the cells in the cytokeratin (CK) channel are "rare" and so at most 3 cells are present in a frame. If this assumption is not met, the parameters need to be changed accordingly to achieve high sensitivity of detection.

It is assumed that the scanned image file names are of the format `Tile%06d.tif`, that 2304 images are generated per channel, and that the different channel images are named sequentially after each other. If differing number of frames are imaged, adjust the frame offsets in the commands accordingly.

1. Run the command below in a cmd prompt window after 20% of the frames are scanned on the CK channel:

   ```
   $ ck_segment <scan_output_dir> <segmentation_dir> <sample_name> \
   1 2304 2305 5 0.995 0.3 3 1
   ```

   where `scan_output_dir` is the path to the directory containing scanned images, `segmentation_dir` is the output directory to store the segmented images, and `sample_name` is an identifier for the sample.

2. Run the command:

   ```
   $ feature_extraction <scan_output_dir> <segmentation_dir> \
   <sample_name> 1 2304 <channel_start>
   ```

   where `channel_start` is a comma separated list of channel start offsets to user for feature extraction. This should be set to `"1, 2305"` when scanned on DAPI and CK channel and to `"2305"` when scanned only on the CK channel.

3. Run the command to filter cells based on feature values:

   ```
   $ filter_features <sample_name>_feature_vec.txt \
   <sample_name>_feature_vec_filt.txt <col(1-based),min,max>*
   ```

   where <col(1-based),min,max>* specifies the column in the feature vector file and the minimum and maximum values in that column to retain. To filter events corresponding to typical cell sizes use `"5,100,10000"`. In addition, filters can also be used for DAPI and CK mean intensities.

4. Run the command to visualize and confirm all the segmented events:

   ```
   $ cell_image <scan_output_dir> <segmentation_dir> \
   <sample_name>_feature_vec_filt.txt "1,1,0,0" "2305,1,0,0" 150
   ```

**Cell relocation to picking:** The cell relocation and picking protocol will vary depending on the make and model of the microscope and micro-manipulator used. We describe a general procedure without specific details.

1. Wipe down all equipment and benches with RNAse away.
2. View the segmented cell events and confirm the cells for picking.
3. Put picking and slide buffers on ice. Add 4µL of RNAse inhibitor to each tube. Vortex briefly.
4. Fill a coplin jar with 1x PBS. Add 4µL of RNAse inhibitor. Mix.
5. To remove slide cover gently and carefully. Place scanned slide in coplin jar and carefully peel off cover.
6. Place slide on the stage of the picking microscope.
7. Add 1000µL of slide buffer to the slide.
8. Find the fiducial point on the slide.
9. The distance to the cell of interest from the fiducial point is calculated using the stage coordinates of the fiducial and that of the frame containing the cell. Move the stage by the calculated distance to go the cell. This process can be automated using a macro.
10. Once the cell is relocated, add 1µL of picking buffer (on ice) to PCR tube. Pick the cell using the micro-manipulator and deposit the cell in the tube. Take the tube off carefully, spin it down, and immediately place it on dry ice.
11. Repeat steps 8-10 until all cells of interest are picked.
12. Store cells in -80°C until needed for RNA extractions.

**dt25 RNA extraction:** The dt25 RNA extraction steps are based on Smart-seq2 (Picelli, 2014) and FRISCR (Thomsen, 2016).

1. Wipe down all equipment with RNAse away, including pipettes, counters, ice bucket, tube holders, and the thermocycler. Prepare buffers at designated time points. Only add RNAse inhibitor 15 minutes before use in each buffer.
2. Get bucket of ice.
3. Place 96-well magnet in -20°C. This will act as a cooler in downstream steps following RNA elution.
4. Turn on thermocycler and prepare the following method.
   (a) 56°C for 1 hour,
   (b) Hold at 56°C (to remove samples and add beads),
   (c) 1 minute hybridization at 56°C (for dt25 beads and polyA transcripts).

5. Prepare RNA lysis buffer in a clean eppendorf tube. The example below is for 12 reactions. Adjust accordingly.
   (a) 8µL of 10x lysis buffer (2% triton X-100; 2x final concentration)
   (b) 2.5µL of Proteinase K (0.0625x final concentration)
   (c) 2.5µL of RNase inhibitor (2U final concentration)
   (d) 27µL of RNase free water (to 40µL)
6. Obtain single cells from -56°C Transport cells from freezer in a 96-well cooler. Once in RNA station, remove tubes from cooler and put in PCR strip.
7. Add 3µL of lysis buffer to each cell.
8. Flick or briefly vortex tubes to mix. Spin down then add tubes to thermocycler.
9. Incubate at 56°C for 1 hour.
10. Place 96-well cooler plate in 4°C. Use this again when preparing final RT rxn.
11. During the hour incubation, prepare buffers, dT25 bead mix, and hybridization mix. Keep buffers and all solutions on ice.
12. Buffers to prepare:
    (a) **2x wash buffer**. To prepare 6mL which is sufficient for 12 reactions:
        i. 1200µL of 20x SSPE (4x final)
        ii. 12µL of 50% tween-20 (0.1% final)
        iii. 3µL of RNase inhibitor (0.05% final)
        iv. 4785µL of RNase free water
    (b) **1x wash buffer**: prepare 1x wash buffer from 2x wash buffer (i.e., take 5mL of 2x wash buffer and 5mL of nuclease free water). Leave 1mL of 2x wash buffer for bead resuspension.
    (c) 10mL **of 1x PBS** in nuclease free water.
    (d) 5mL **of elute buffer** – nuclease free water.
13. **dt25 bead preparation**: remove 8µL of dt25 beads for each sample and add to clean 200µL PCR or 1.5mL eppendorf tube (depending on volume).
    (a) Add dt25 beads to magnet. Wait 2 minutes. Remove supernatant and add 200µL of 1x wash buffer. Vortex. Spin down briefly. Be sure to resuspend the beads so they are washed thoroughly.
    (b) Repeat 200µL of 1x wash 2 more times (3 total times).
    (c) Following the third wash, return beads to magnet. Wait 2 minutes. Remove 200µL. Pipette all liquid out of the tube while beads are on magnet.
    (d) Add 4µL of 2x wash buffer to beads for each sample (half volume of beads intially taken).
    (e) Keep beads on ice or at 4°C until ready.
    (f) For example, if you have 100 samples, you will use 800µL of beads and then resuspend the washed beads in 400µL of 2x wash buffer.

14. **Primer hybridization mix preparation**:
    (a) Remove dNTP mix from -20°C and the dT primer from -80°C.
    (b) For each reaction, add 0.5µL of dNTP mix and 1µL of dT primer (with about 20% extra solution). For example, for 32 samples, add 40µL of dT primer and 20µL of dNTP mix.
    (c) In clean PCR strips, add 1.5µL of primer hyb mix to each tube. This is where RNA will be eluted from dT25 beads and added to.
    (d) Label and strips in 4°C until elution.
15. Prepare tubes for gDNA supernatant in PCR strips if you wish to save for WGA. Set these aside for bead purification.
16. Following 1 hour incubation, remove samples from incubator and place in regular 96-well PCR strip. Keep at room temperature.
17. Take beads and mix so there is no sedimentation at bottom of bead tube.
18. Add 4µL of beads to each sample. Flick tubes and briefly spin down.
19. Incubate at 56°C for 1 minute.
20. Remove from thermocycler and place tubes at room temperature on RNA bench for 5 minutes.
21. Add 5µL of RNAse inhibitor to 1x PBS buffer and Elute buffer. 1X wash buffer has RNAse inhibitor from 2x wash buffer.
22. Place samples with beads in them on room temp 96 strip magnet for 1 minute.
23. Carefully, without disturbing beads, remove supernatant with 10µL multichannel pipette and add supernatant to prepared tubes for gDNA. You should have 5µL of supernatant. Store supernatant for WGA at 4°C until RNA extraction is complete.
    (a) Alternatively, if you do not wish to profile the genome, throw away supernant and continue with washing the beads.
24. Keeping beads on magnet, wash beads twice with 100µL of 1x wash buffer. Add in buffer, let sit for 30 seconds, then remove and add again. Make sure beads are not disturbed.
25. Remove 1x wash buffer and add 100µL of 1x PBS.
26. With pipette set to 150µL, remove 1x PBS. Try to remove all liquid from tubes.
27. Remove tubes from magnet and place in regular 96 tube PCR strip.
28. Add 3.5µL of elute buffer (water + RNase inhibitor). Flick tubes to re-suspend beads. Briefly spin down tubes.
29. Elute RNA from beads by incubating at 80°C for 20 mins.
30. During this time, (1) remove primer hybridization mix from 4°C and (2) remove 96-well magnetic plate from -20°C.

31. Remove tubes from thermocycler and place in ice cold magnetic stand.

32. Set thermocyler to 72°C hybridization step (72°C for 3 minutes and 4°C for 3 minutes).

33. Using pipette set to 7µL, transfer supernatant to tubes containing primer hybridization mix. Vortex and spin down. Add to thermocylcer and incubate for 72°C for 3 minutes and 4°C for 3 minutes.

34. Prepare RT reaction mixture. Example below is for 1 sample.

    (a) 2µL 5x first strand buffer
    (b) 0.5µL TSO (template switching oligo)
    (c) 0.25µL RNase inhibitor
    (d) 1µL SMARTScribe Reverse Transcriptase

35. Using cooler stored in 4°C, remove tubes from thermocycler. Add 3.75µL of RT reaction mixture. Flick to vortex and briefly spin down.

36. Set thermocycler to RT reaction program:

    - Incubate at 42°C for 90 minutes to allow for RT and template switching
    - Cycle 10 rounds of:
      - 50°C for 2 minutes to unfold RNA secondary structures.
      - 42°C for 2 minutes for completion and continuation of RT and template switching.
    - Inactivate enzyme at 70°C for 15 minutes.
    - Hold at 4°C

37. Add tubes to thermocycler.

38. cDNA can be stored overnight at 4°C or at -20°C for at least 1 month, however it is easiest to move forward with cDNA amplification.

39. If you decide to proceed with WGA on the gDNA fraction, proceed to GeTMoR DNA WGA **immediately**. Since the gDNA fraction of a cell is isolated and lysed, WGA needs to be conducted on the same day as RT extraction or DNA will be lost.

**cDNA amplification:** This step immediately follows RT conversion to cDNA (step above). It's easiest and suggested to perform immediately after the RT reaction.

1. Remove tubes from incubator or storage.

2. To each tube, add 15µL of the following master mix solution

    - 12.5µL of 2x KAPA mix
    - 0.25µL of ISPCR
    - 2.25µL of nuclease free water

3. Perform PCR using the KAPA program

- Denature for 1 round at 98°C for 3 minutes
- Cycle 21 rounds of:
  - Denature at 98°C for 20 seconds
  - Anneal at 67°C for 15 seconds
  - Extend at 72°C for 6 minutes
- Final extension for 5 minutes at 72°C
- Hold at 4°C

4. Tubes can be stored overnight at 4°C or at -20°C for at least 1 month, but it is suggested to purify with XP PURE beads before storage.

**cDNA purification:**

1. Equilibrate beads at room temp for 30 minutes prior to use.
2. Add 0.8X beads per reaction – add 15μL of room temp XP beads to each amplified cDNA reaction.
3. Incubate at room temperature for 8 minutes.
4. Add beads to magnetic stand for 5 minutes or until solution is completely clear of bead mix.
5. Remove supernatant and discard.
6. Add 200μL of freshly prepared 80% ethanol. Incubate for 30 seconds. And remove.
7. Repeat step 6 (above).
8. Remove all liquid from tube and incubate on magnet for 3 minutes (DO NOT OVER-DRY BEADS).
9. Add 14μL of RNase free water. Vortex vigorously. Pat tubes on counter to try and make sure beads settle towards bottom.
10. Incubate for 5 minutes. Then spin down tubes and place on magnet for 3 minutes.
11. Elute 12μLof purified cDNA into clean PCR tubes.
12. Quantify cDNA yield with quibit (2μL sample) and cDNA integrity with bioanalyzer (1μL sample). If cells pass QC, proceed to library prep via Nextera XT.

**Nextera XT cDNA library preparation (1/5 reactions):**

1. Dilute cDNA to 250 pg (0.250 ng) total into PCR strips with pure water. Make minimum of 5μL of diluted cDNA.
2. Prepare transposase master mix and 3μL of mix to each tube:
   (a) 2μL transposase buffer
   (b) 1μL transposase
3. Set thermocycler to the following:

(a) 56°C hold
(b) 56°C for 10 minutes
(c) 10°C hold

4. Add 1µL of diluted cDNA to transposase mix. Vortex briefly, spin down, and place in thermocycler and start 56°C incubation.
5. Prepare stop NT buffer and get ready to add to samples when they are finished with transposase reaction.
6. Remove tubes and add 1µL of stop buffer. Vortex and spin down.
7. Wait 5 minutes.
8. Add 3µL of NPM pcr mix.
9. Add 1µL of each i5 and i7 PCR primer.
10. Vortex and spin down.
11. Perform PCR using the nextera PCR program
    - Gap fill for 1 round for 3 minutes at 72°C
    - Initial 95°C denaturation for 30 seconds
    - Cycle 12 rounds of:
        - Denature at 95°C for 10 seconds
        - Anneal at 55°C for 30 seconds
        - Extend at 72°C for 1 minute
    - Final extension for 5 minutes at 72°C
    - Hold at 4°C. Either proceed to purification within 24 hours, or store at -20°C for up to 1 month.
12. Purify with 0.8X XP Pure Beads

**GeTMoR DNA WGA:** WGA is meant to take place on the same day as RNA extractions; the gDNA cannot be stored or else WGA will fail. The RNA lysis step with proteinase K also lyses the nucleus and releases gDNA – this step takes over the DTT-KOH lysis for SIGMA WGA4 protocol. You will begin by adding in the fragmentation mix (10x mix) from the WGA4 protocol but adjust for volume discrepancies. It is assumed that you have roughly 5μL of gDNA in each tube.

1. Prepare 2x fragmentation mix: 1μL 10x fragmentation + 4μLTE buffer.
2. Add 5μL of 2x fragmentation mix to each sample. Mix by vortexing and spin down.
3. Incubate at 99°C for 4 minutes.
4. Remove samples and place in new PCR tube cooler from genomics -20°C.
5. Make library preparation master mix
   (a) 2μLof 1x single cell library preparation buffer (green cap)
   (b) 1μL of library stabilization solution (yellow cap).
6. Add 3μL of library prep master mix to each sample. Mix and place in thermocycler at 95°C for 2 minutes. Cool samples on PCR tube cooler.
7. Add 1μL of library preparation enzyme (orange cap), mix by flicking tube (enzyme is fragile), and spin down.
8. Place sample in thermocycler and incubate as follows:
   (a) 16°C for 20 minutes
   (b) 24°C for 20 minutes
   (c) 37°C for 20 minutes
   (d) 75°C for 5 minutes
   (e) Hold reactions at 4°C
9. Make amplification master mix
   (a) 7.5μL of 10x amplification master mix (red cap)
   (b) 48.5μL of molecular grade water
   (c) 48.5μL of WGA DNA polymerase (white cap)
10. Add 61μL of master mix to each reaction. Mix by vortexing and spin down.
11. PCR reaction of:
    (a) Initial denaturation at 95°C for 3 minutes
    (b) 23 cycles of
        i. 94°C denature for 30 seconds
        ii. 65°C anneal/extend for 5 minutes
    (c) Hold at 4°C
12. Follow WGA4 quality control metrics: (1) run gel and look for smearing pattern with minimal primer dimers. If gel looks good, then (2) purify samples with spin columns.
